# Do microsaccades vary with discriminability around the visual field?

**DOI:** 10.1101/2024.01.11.575288

**Authors:** Simran Purokayastha, Mariel Roberts, Marisa Carrasco

## Abstract

Microsaccades–tiny fixational eye movements– improve discriminability in high acuity tasks in the foveola. To investigate whether they help compensate for low discriminability at perifovea, we examined MS characteristics relative to the adult visual performance field, which is characterized by two perceptual asymmetries: Horizontal-Vertical Anisotropy (better discrimination along the horizontal than vertical meridian), and Vertical Meridian Asymmetry (better discrimination along the lower-than upper-vertical meridian). We investigated whether and to what extent microsaccade directionality varies when stimuli are at isoeccentric locations along the cardinals under conditions of heterogeneous discriminability (Experiment 1) and homogeneous discriminability, equated by adjusting stimulus contrast (Experiment 2). Participants performed a two-alternative forced-choice orientation discrimination task. In both experiments, performance was better on trials without microsaccades between ready signal onset and stimulus offset than on trials with microsaccades. Across the trial sequence the microsaccade rate and directional pattern were similar across locations. Our results indicate that microsaccades were similar regardless of stimulus discriminability and target location, except during the response period–once the stimuli were no longer present and target location no longer uncertain–when microsaccades were biased toward the target location. Thus, this study reveals that microsaccades do not flexibly adapt as a function of varying discriminability in a basic visual task around the visual field.

## Introduction

### Behavioral performance fields

Our vision is not uniform across the visual field: we see most clearly directly ahead (i.e., where our gaze is) and less clearly toward the periphery. Discriminability and the speed of information processing differ both as a function of eccentricity (reviews: Strasburger et al., 2011; Anton-Erxleben & Carrasco, 2013; Carrasco & Barbot, 2014) and as a function of polar angle, i.e., around the visual field (review: Himmelberg, Winawer & Carrasco, 2023). The pattern of visual performance as a function of polar angle at isoeccentric locations, known as a *performance field* (PF), has been well-documented for several fundamental visual dimensions, e.g., contrast sensitivity (e.g., Rovamo & Virsu, 1979; Rijsdijk et al., 1980; Carrasco et al., 2001; Cameron et al., 2002; Baldwin et al., 2012; Himmelberg et al., 2020) and appearance (Fuller et al., 2008), spatial resolution (e.g., Nazir, 1992; Altpeter et al., 2000; Talgar & Carrasco, 2002; Montaser-Khousari & Carrasco, 2009; Greenwood et al., 2017; Barbot et al., 2021), motion (Fuller & Carrasco, 2009), spatial crowding (Petrov & Meleshkevich, 2011; Fortenbaugh et al., 2015; Greenwood et al., 2017; Kurzawski et al., 2023), and temporal information accrual (Carrasco et al., 2004). **Figure 1** shows a schematic representation of the typical PF shape for adults.

**Figure 1.**
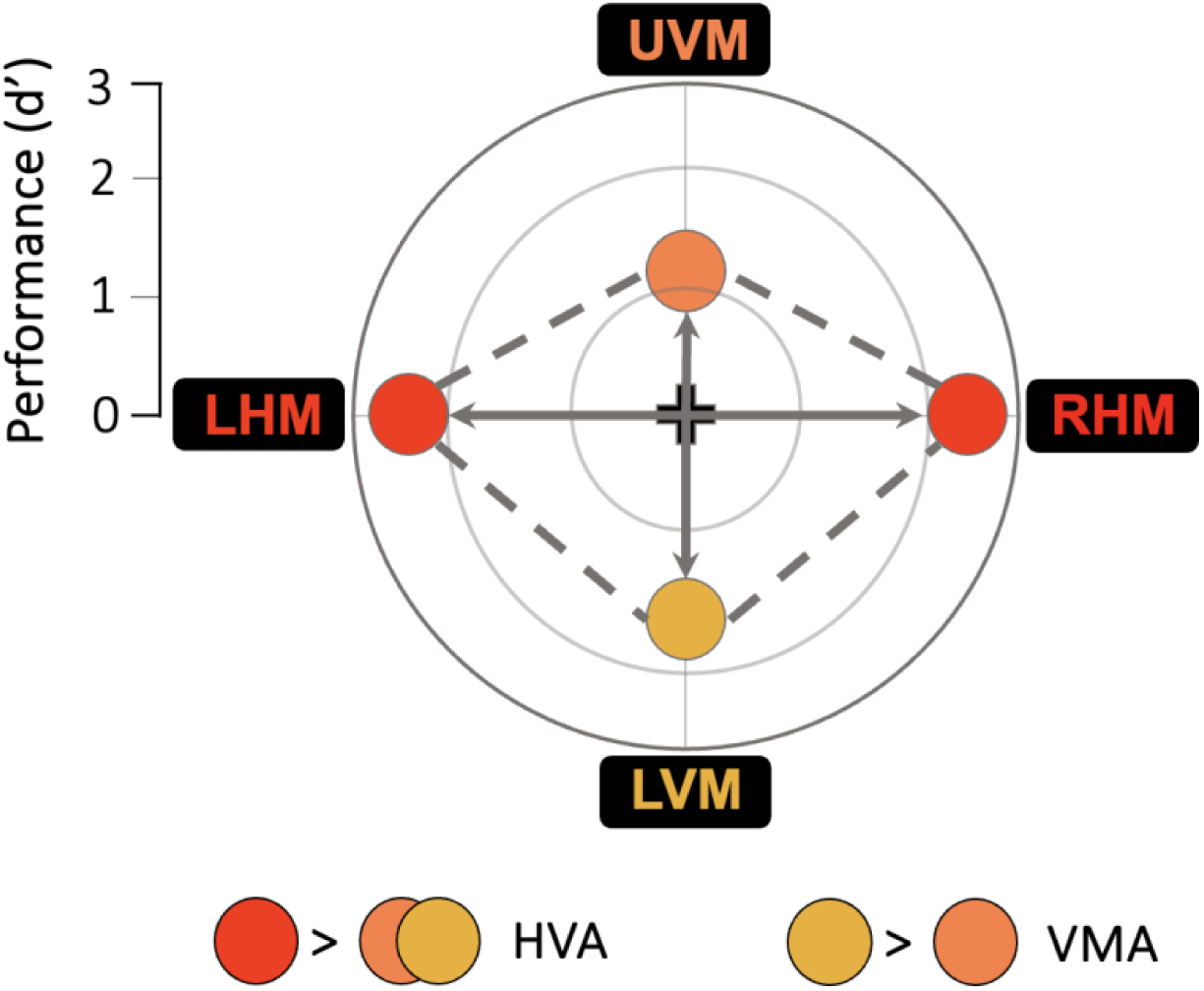
Schematic representation of a typical visual performance field. The center point (intersection of the cardinal lines) represents chance performance, and more eccentric points represent higher performance around the visual field. (1) Horizontal-vertical anisotropy (HVA): better performance at isoeccentric locations along the horizontal (left horizontal meridian (LHM) and right horizontal meridian (RHM) locations) than vertical (upper vertical meridian (UVM) and lower vertical meridian (LVM) locations) meridian of the visual field, and (2) vertical meridian asymmetry (VMA), better performance at the location directly below fixation (LVM) than directly above (UVM).

Adult PFs are characterized by two asymmetries: the HVA (*Horizontal-Vertical Anisotropy*) and VMA (*Vertical Meridian Asymmetry*). The HVA reflects better performance along the horizontal meridian (HM) than the vertical meridian (VM). The VMA refers to better performance along the lower vertical meridian (LVM) than along the upper vertical meridian (UVM). Typically, performance at intercardinal locations is intermediate between that at the horizontal and vertical meridians (e.g., Altpeter et al., 2000; Carrasco et al., 2001; Cameron et al., 2002; Baldwin et al., 2012; Corbett & Carrasco, 2011), and both asymmetries decrease gradually as the target location shifts from the vertical meridian toward the horizontal meridian (Abrams et al., 2012; Barbot et al., 2021).

The HVA and VMA are pervasive; they emerge regardless of stimulus orientation, display luminance, and whether the stimuli are presented monocularly or binocularly, and with or without location uncertainty (Carrasco et al., 2001; Corbett & Carrasco, 2011; Abrams et al., 2012; Purokayastha et al., 2021; Barbot et al., 2021). PFs shift in line with egocentric referents, corresponding to the stimulus retinal location (Corbett & Carrasco, 2011). PF asymmetries are more pronounced as spatial frequency, eccentricity, or set size increase (Rijsdijk et al., 1980; Carrasco et al., 2001; Cameron et al., 2002; Fuller et al., 2008; Baldwin et al., 2012; Greenwood et al., 2017; Himmelberg et al., 2020). As a result of these asymmetries, we naturally perceive objects at different levels of discriminability in daily life, depending on their position in our visual field.

### Microsaccades

Acute vision is possible because of the ability of our eyes to align with a visual target. We do this – voluntarily or involuntarily – using two types of eye movements: those that direct our line of sight to a new object of interest and those that stabilize gaze. Our eyes move continuously even during fixation (reviews: Rolfs, 2009; Martinez-Conde et al., 2013; Rucci & Poletti, 2015). Microsaccades (MS) are one such fixational eye movement: they are small (< 1° of visual angle), often occur involuntarily, and are typically conjugate (Møller et al., 2002; Martinez-Conde et al., 2004; Collewijn & Kowler, 2008; Rucci & Poletti, 2015). They prevent perceptual fading (e.g., Tulunay-Keesey, 1982; Martinez-Conde, 2006; McCamy et al., 2012), are informative oculomotor correlates to perception (Dankner et al., 2017; Fried et al., 2014; Gooding & Basso, 2008; Rucci & Poletti, 2015), and have been functionally linked to the improved detection of fine spatial detail by properly relocating the gaze in high-acuity tasks within the parafoveal region (Ko et al., 2010), and even within the small foveal subregion called the foveola (e.g., Rucci et al., 2007; Donner & Hemilä, 2007; Ko et al., 2010; Poletti et al., 2013; Rucci & Poletti, 2015).

Microsaccade dynamics – including changes in rate and direction – are also associated with a variety of cognitive functions, including working memory (van Ede et al., 2019; Valsecchi et al., 2007; Martinez-Conde et al., 2009; Valsecchi & Turatto, 2009; Willeke et al., 2019), perceptual learning (Hung et al., 2022, 2023), temporal expectation (Dankner et al., 2017; Amit et al., 2019; Abeles et al., 2020; Badde et al., 2020), temporal attention (Denison, Yuval-Greenberg & Carrasco, 2019; Palmieri, Fernández & Carrasco, 2023) and spatial attention (although their role is controversial; Hafed & Clark, 2002; Engbert & Kliegl, 2003; Yuval-Greenberg et al., 2014; Lowet et al., 2018; Roberts & Carrasco, 2019; Xue et al., 2020; Yu et al., 2022).

Studies in which observers were required to maintain central fixation have reported that MS direction is biased along the horizontal meridian, with much fewer MS to the vertical meridian, and very few to oblique directions (Hafed & Clark, 2002; Engbert & Kliegl, 2003; Laubrock et al., 2005; Liang et al., 2005; Turatto et al., 2007; Yokoyama et al., 2012; Yuval-Greenberg et al., 2014; Raveendran et al., 2020). However, it is important to note that most of these studies only placed stimuli along the horizontal meridian. Of the studies that placed stimuli at locations other than on the horizontal meridian, the experimenters only tracked and analyzed horizontal MS (Hafed & Clark, 2002), used microsaccade-contingent stimuli- and cue- presentation (Yuval-Greenberg et al., 2014), or used a visual search task with high location uncertainty (Turatto et al., 2007).

### Microsaccades and performance asymmetries

Performance asymmetries around the visual field are well-documented, but only one study on temporal attention has analyzed MS dynamics as a function of polar angle, ‘around’ the visual field (Palmieri et al., 2023). Knowing that behavioral PFs reflect differences in discriminability, in this study we investigated whether and to what extent the directionality of MS varies when stimuli are placed at isoeccentric locations along the cardinals. Do they parallel behavioral PF patterns (i.e., are biased toward the locations with higher discriminability), compensate for them (i.e., are biased toward the locations with lower discriminability), or seem independent of target location? Observers performed a 2AFC orientation discrimination task under two conditions of varying discriminability around the visual field. In one condition –heterogeneous discriminability– the stimulus contrast was the same at all locations, and thus discriminability across isoeccentric locations varied, representative of real-world conditions (Experiment 1). In the other condition –homogeneous discriminability– stimulus contrast at each location was adjusted to equate discriminability, so that locations were at the same baseline level (Experiment 2) enabling us to document potential directional biases in MS independent of varying discriminability. This way, if MS differ around the visual field, we could disentangle whether the distinct patterns would reflect differences in discriminability as a function of the current task demand or typical differences in discriminability around the visual field.

## Methods

We employed a standard two-alternative forced-choice (2AFC) orientation discrimination task to investigate the relation between MS and canonical PF asymmetries. We performed MS analyses on the neutral condition of a published data set (Purokayastha et al., 2021) and on new data from eight additional observers. We collected data under two conditions: heterogeneous discriminability (Experiment 1) and homogeneous discriminability (Experiment 2). The procedure from the published experiment is summarized here. New analyses implemented in the current paper are described in detail.

### Participants

The same 28 adults (23 female; mean age = 24.21 ± 4.76 years) participated in both experiments, all of whom possessed normal- or corrected-to-normal vision and were attending college or graduate school at New York University (NYU). All experimental procedures were in agreement with the Declaration of Helsinki. The protocols for the study were approved by the NYU Institutional Review Board. We obtained informed consent from all participants.

### Apparatus and setup

Participants were tested in a dimly lit, sound-attenuated room. The stimuli were programmed on an Apple iMac MC413LL/A 21.5-in. Desktop (3.06 GHz Intel Core 2 Duo) using MATLAB (Mathworks, Natick, Massachusetts, USA) in conjunction with the MGL toolbox (http://gru.stanford.edu/doku.php/mgl/overview). Stimuli were presented at a viewing distance of 57cm on a 21-in. IBM P260 CRT monitor (1280x960 pixel resolution, 90-Hz refresh rate), calibrated and linearized using a Photo Research (Chatworth, CA) PR-650 SpectraScan Colorimeter.

Participants performed the task using a forehead- and chin- rest to minimize head movements. Eye movements were monocularly monitored (right eye) using an EyeLink 1000 Desktop Mount eye tracker (SR Research, Ontario, Canada) with a spatial resolution of 0.01dva and a temporal resolution of 500Hz. A standard 5-point grid calibration was used every 4 blocks or when there were many fixation breaks in a given block.

### Stimuli

The stimuli were the same across all sessions except for the contrast of the Gabor patch stimuli (see Procedure below). Observers were asked to fixate on a black central cross (0.5° across) throughout the trial (**Figure 2**). Four placeholders—each composed of four black dots (0.05° radii) arranged in a circle 0.5° from the edge of an upcoming Gabor patch stimulus (to prevent masking)—were always presented on the screen to reduce location uncertainty. The target and three distractor stimuli were 3.2° wide Gabor patches (contrast-defined sinusoidal gratings embedded in a Gaussian envelope, σ = 0.46°) with a spatial frequency of 4 cycles per degree (cpd), and the same mean luminance as the uniform gray background (26 cd/m^2^). They were centered at 6.4° eccentricity along the cardinal meridians. During each trial, the Gabors were independently and randomly tilted ±20° from vertical. The ready signal consisted of four 0.28° lines (all 0.14° thick)—0.38° from the center of the fixation cross, indicating all four possible target locations. A response cue (a single 0.88° line, 0.38° from the center of the fixation cross) indicated the target location by pointing to one placeholder and eliminated location uncertainty at response time (Pelli, 1985; Carrasco et al., 2002).

**Figure 2.**
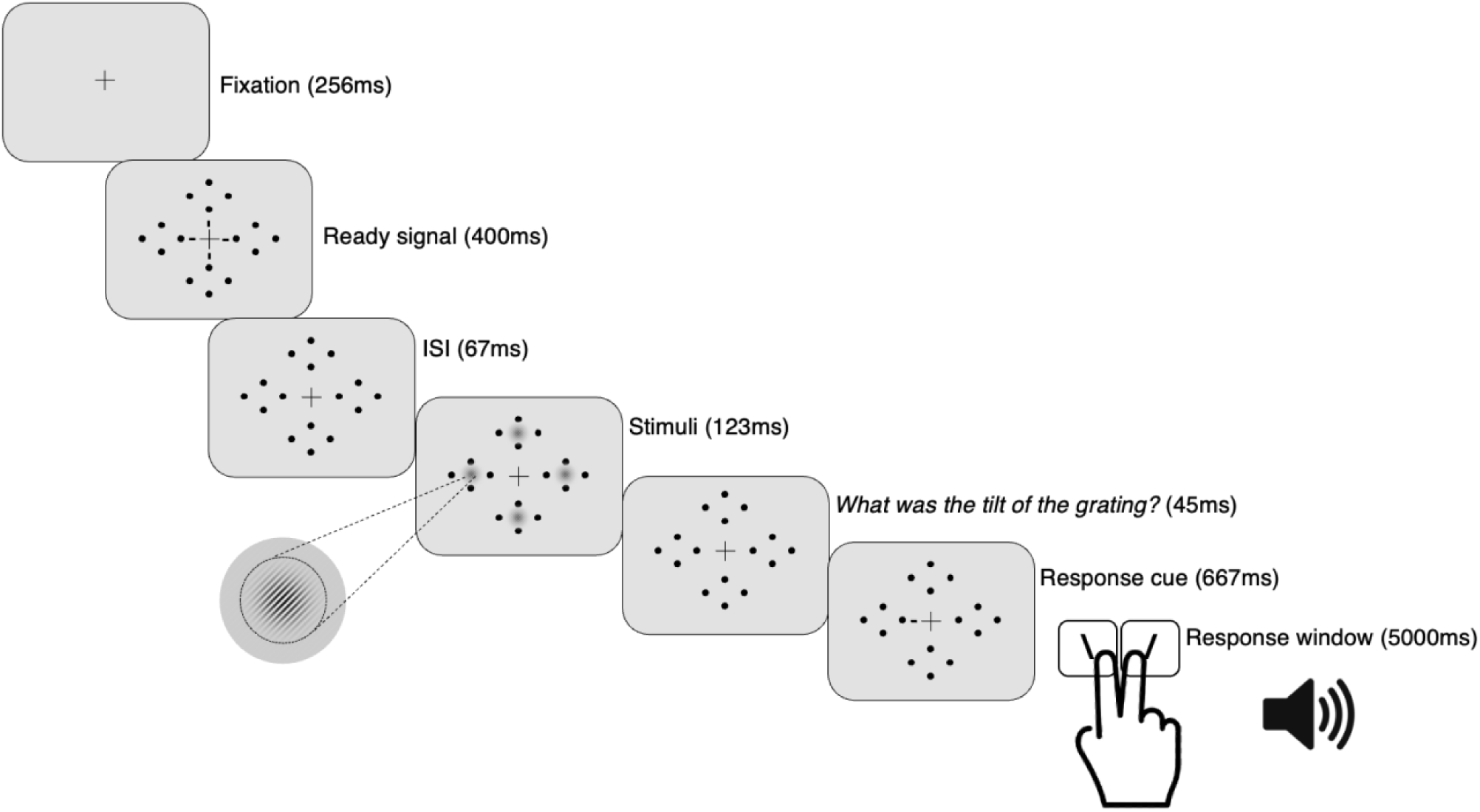
Task sequence. (**a**) Observers performed a two-alternative forced-choice (2AFC) orientation discrimination task – contingent on contrast sensitivity – while maintaining fixation. The target could appear at any one of four isoeccentric locations along the cardinals. On every trial, four Gabor patches briefly appeared simultaneously at all stimulus locations. A response cue indicated the target, for which participants reported the tilt (left or right). Note that Gabor tilt angle, size of placeholders, fixation point, and response cue have been exaggerated for clarity.

### Procedure

Participants underwent three sessions: threshold estimation, heterogeneous discriminability, and homogeneous discriminability. In the first session, we measured observers’ individual stimulus contrast thresholds (8 blocks, ∼1 hour), using a PEST staircase procedure in which eight interleaved 3-down, 1-up staircases (2 staircases per location; one starting from 75% and the other from 25% contrast) estimated a single contrast threshold value (∼80% accuracy overall) for each location; observers were asked to report the orientation of a single target without any distractors on each trial. Next, the order of the heterogeneous- and homogeneous- discriminability experiments was counterbalanced across participants. Both 45-min sessions consisted of 12 blocks of 40 trials each, for a total of 480 trials (120 trials for each location, presented in random order).

Experiment 1: In the <u>heterogeneous</u> discriminability across locations experiment, Gabors at all locations were presented at the same contrast, which was the average of the contrast thresholds independently measured at the four potential target locations (**Figure 3a**). In between blocks, stimulus contrast was adjusted to maintain ∼80% accuracy across all locations and cueing conditions.

**Figure 3.**
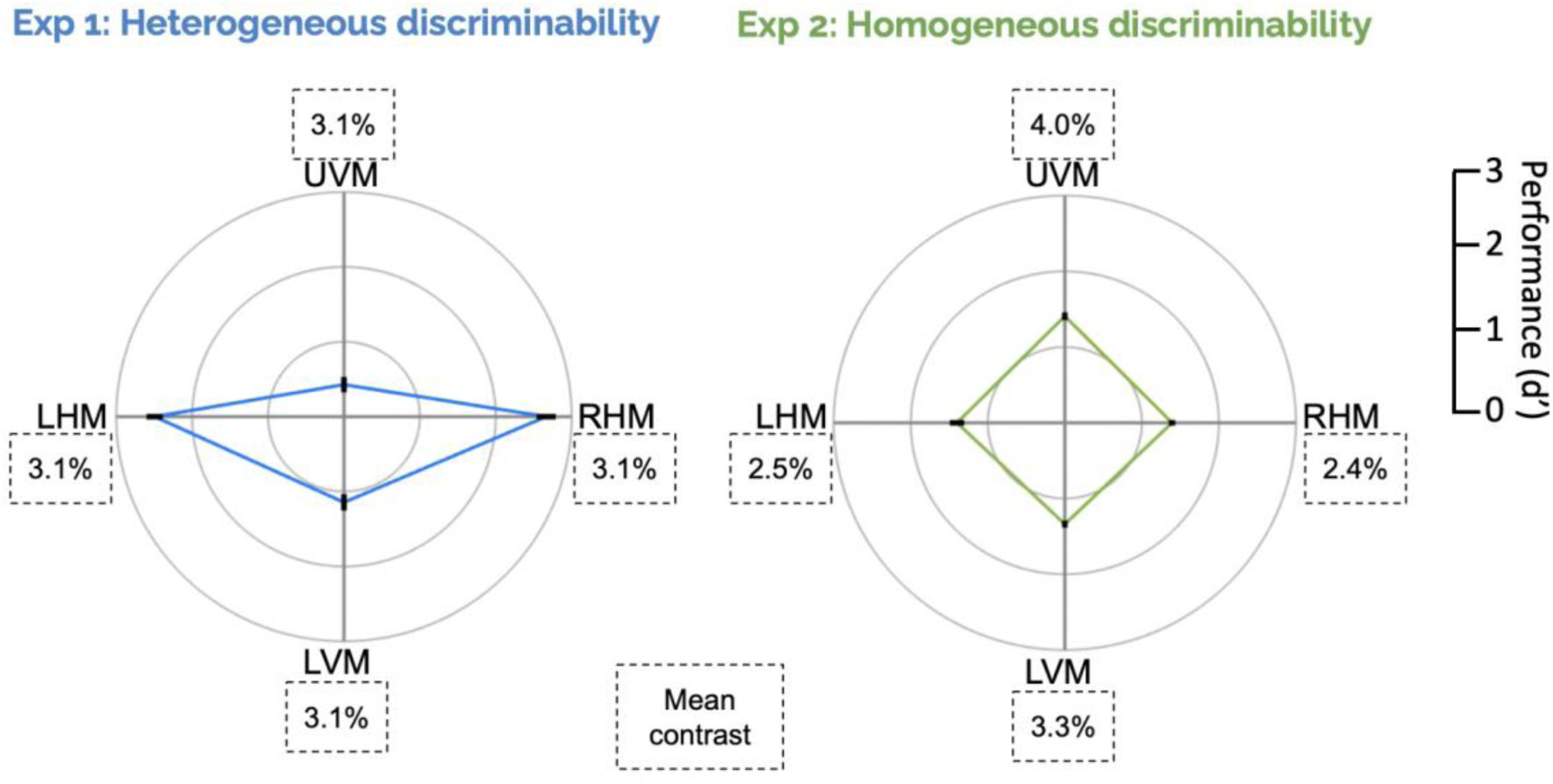
Task performance in both experiments. Group mean stimulus contrast by location is shown in dotted-line boxes for each experiment while the polar plots show mean d’ performance at all tested locations, with black lines representing ±1 SEM. In Experiment 1 (left polar plot), the same contrast across locations resulted in different performance across locations; in Experiment 2 (right polar plot), stimuli were presented at a different contrast at each location to equate performance (∼80% accuracy) across locations.

Experiment 2: In the <u>homogeneous</u> discriminability across locations experiment, we equated discriminability across the visual field by presenting stimuli of different contrasts according to the independently measured contrast thresholds of each location. In between blocks, stimulus contrast was adjusted separately for each location to maintain 80% accuracy across cueing conditions (**Figure 3a**). Doing so ensured that participants performed well above chance and had similar room for improvement at all locations.

Participants were encouraged to take a short rest in between blocks if they felt fatigued. Throughout all sessions, if participants made an eye movement outside of a 1° radius from the fixation cross at any time between trial initiation and stimulus offset, the trial was aborted and the text, ‘Please fixate’, would appear at the center of the screen. These trials (Experiment 1: 6.17%, Experiment 2: 4.96%) were rerun at the end of each block.

### Task and trial sequence

In both experiments, participants performed a 2AFC orientation discrimination task binocularly (**Figure 2**). Once observers had fixated for 256ms, the ready signal was presented for 400ms, followed by a brief blank stimulus onset asynchrony (SOA) of 67ms. After the blank interval, four Gabor patches appeared simultaneously inside the placeholders for 123ms. There was a brief 45- ms interstimulus interval (ISI) between display offset and the response cue, which was presented for 667ms. A mid-frequency auditory tone indicated the beginning of the response window in which observers had to report the target tilt (clockwise (CW) or counterclockwise (CCW) relative to vertical) using one of two keyboard presses (1 for CCW, 2 for CW) with their right hand. On every trial, observers were encouraged to respond as accurately as possible. Observer response terminated the response window, after which there was a mandatory 1-s intertrial interval. Auditory feedback was provided at the end of each trial (low-frequency tone: incorrect; high-frequency tone: correct) and visual feedback indicating observers’ accuracy and number of fixation breaks was presented onscreen at the end of each block.

### Analysis

We used an alpha level of 0.05 for all statistical tests.

### Behavior

We calculated d’ as the z-score of the proportion of hits (target was CW, observer reported CW) minus the z-score of the proportion of false alarms (target was CCW, observer reported CW) (e.g., Herrmann et al., 2010; Zhang et al., 2019; Jigo & Carrasco, 2020). To avoid infinite values, we adjusted all d’ values using the conservative log-linear rule by adding 0.5 to the number of hits and false alarms before computing d’ (Hautus, 1995).

### Microsaccades

#### Microsaccade detection

All eye tracker data were preprocessed using the Engbert and Kleigl (2003) velocity-based detection algorithm to identify MS; detection thresholds were determined in two-dimensional (2D) velocity space computed separately for horizontal and vertical components. The raw time series of eye positions were transformed into 2D velocity space with a threshold of at least 6*SD* above the average velocity per trial for at least 6 consecutive time points. MS were defined as saccades with an amplitude smaller than 1dva, a duration between 8ms and 40ms, and a peak velocity less than 100dva/s. We imposed a minimum intersaccadic interval (defined as the interval between the last sample of one saccade and the first sample of the next saccade) of 10ms so that potential overshoot corrections were not considered new MS. The algorithm is robust with respect to different noise levels between different trials and observers.

#### Relation between microsaccades and task performance

To link MS dynamics to established behavioral PF asymmetries, we compared task performance on trials in which there was at least one MS during a critical period for performance (between ready signal onset and stimulus offset) versus trials without MS across all observers using a Bonferroni-corrected paired t-test. To obtain a measure of performance, we calculated proportions of hits and false alarms for each block and for every observer, summing them per observer and calculating d’ as detailed above. We also compared mean reaction times (from response cue onset) across blocks for every observer in trials with- and without- MS, to rule out the possibility of a speed-accuracy trade-off.

#### Microsaccade rate

The microsaccade rate (MS/s) for the entire trial sequence was calculated continuously as the total number of MS divided by the total number of trials per second for that observer on each trial. All MS rate data were smoothed using a moving mean with a 50-ms Gaussian sliding window filter. We examined MS rate across the entire trial sequence –for both experiments and split by correct and incorrect trials– all locked to ready signal onset.

To determine whether any timeseries differences were statistically significant, we used a nonparametric cluster-based permutation test, which determines whether any observed cluster of time points is significantly different from that expected by chance (Maris & Oostenveld, 2007). In implementing this test, we used the largest absolute summed t-values within a cluster, shuffled the experiment/behavioral response accuracy labels for each observer, and repeated the process 1000 times. The output from each iteration was the largest cluster from that set, and together, these formed a null distribution of clusters. A cluster was determined to be statistically significant if it was greater than 95% in the null distribution (*p* < 0.05).

#### Microsaccade direction

In analyzing the spatial distribution of MS throughout the trial, we divided the polar field into 16 evenly sized (22.5°-wide) segments and calculated the MS frequency in the direction of each segment over the full trial sequence. To assess how MS directionality differed during this time period, we used Bonferroni-corrected paired t-tests to test the null hypothesis that the number of MS that occurred toward the three central wedges (67.5°) in each cardinal direction (right HM (RHM), UVM, left HM (LHM), and LVM) were all equal (*p* < .05) (**Figure 7a**).

#### Gaze position

We also examined the distribution of gaze positions around the visual field to determine whether it was biased in any way and could be related to MS direction. To do this, we used gaze position data collected by the eye tracker (in Cartesian coordinates) for every trial, converted them to polar coordinates, and aligned them with respect to the screen center. We then averaged the resulting gaze data per trial for every observer to plot the spread of gaze positions around the visual field across the full sequence and response cue period. We also calculated the ‘grand average gaze position’ as the average gaze position across all trials and all observers, represented relative to screen center.

## Results

### Behavior

Detailed behavioral results are provided in the neutral conditions of Purokayastha et al. (2021) and summarized here (including 20 from published study and 8 more observers). In Experiment 1, when stimuli at all locations had the same contrast (*M* contrast=3.1%, 95% CI [2.6%, 3.6%]), observers exhibited canonical performance fields (**Figure 3, left panel**); participants showed pronounced and consistent HVA, i.e., higher performance accuracy along the HM than the VM, and VMA, i.e., higher accuracy at the LVM than the UVM. In Experiment 2, we equated stimulus discriminability by adjusting stimulus contrast (**Figure 3, right panel**), ensuring that all locations were reliably above chance and below ceiling. Our contrast manipulation was successful; performance was equated across all locations around the visual field. We also analyzed reaction time data in both experiments and confirmed that there were no speed-accuracy tradeoffs.

### Microsaccades

We discarded 5.95% of the data (Experiment 1: 6.02%, Experiment 2: 5.88%) after preprocessing (trials without a response, or when gaze position drifted > 1dva from the central fixation cross). The total number of MS detected across all participants and sessions and used for final analyses was 38,867 (Experiment 1: 19,939, Experiment 2: 18,928).

### Main sequence and related microsaccade dynamics

Converging research indicates that MS and saccades are generated by the same underlying circuitry (Martinez-Conde, 2006; Otero-Millan et al., 2008; Mergenthaler & Engbert, 2010; Poletti & Rucci, 2016). Accordingly, saccades and MS identified by the algorithm followed the saccadic main sequence (Zuber et al., 1965), i.e., MS amplitudes and peak velocities were highly correlated, Experiment 1: Pearson’s *r* = .9567, *p* < .001, 95% CI [0.9562, 0.9572], Experiment 2: Pearson’s *r* = .9610, *p* < .001, CI [0.9606, 0.9615] (**Figure 4**).

**Figure 4.**
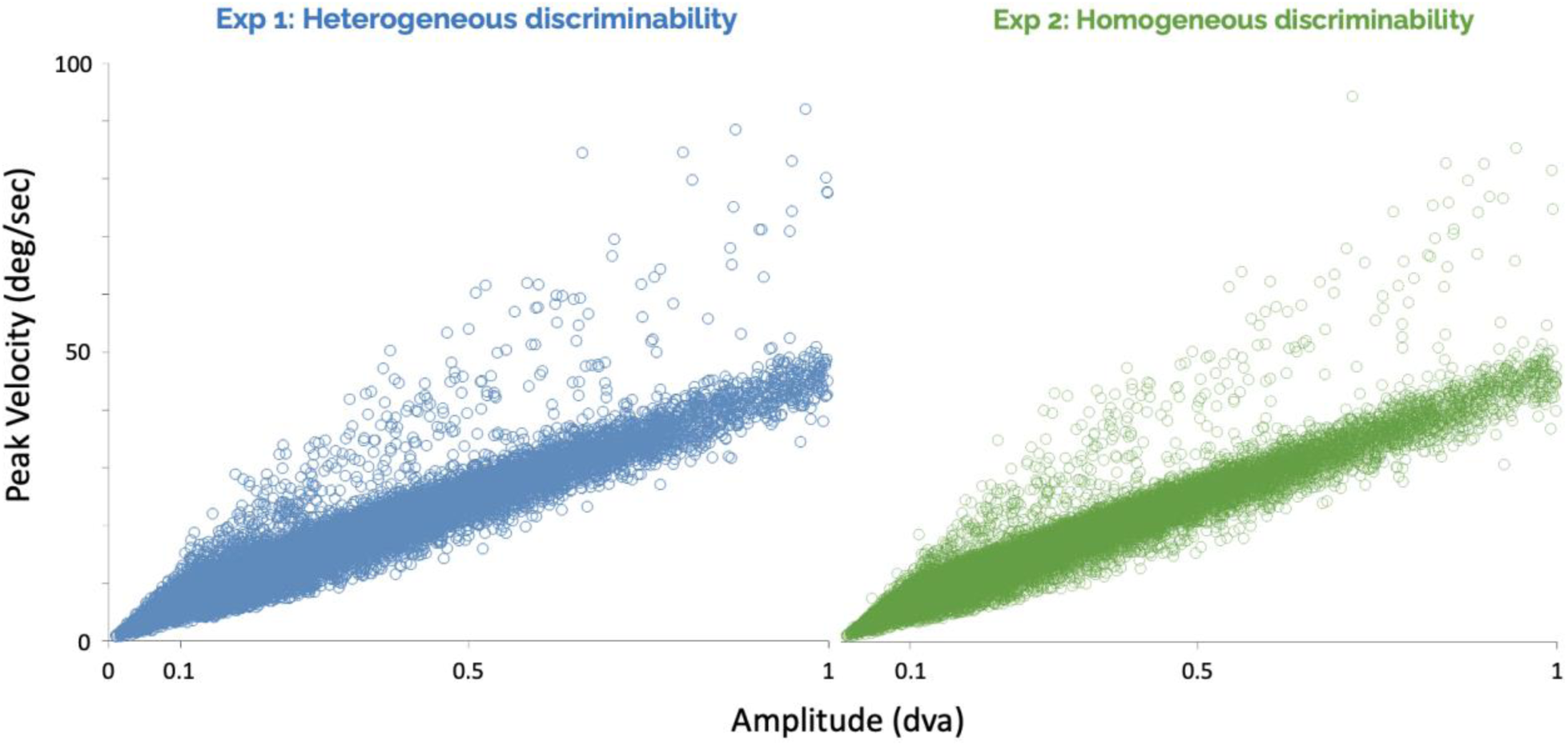
MS peak velocity plotted against MS amplitude for Experiment 1 (heterogeneous discriminability; left panel) and Experiment 2 (homogeneous discriminability; right panel). MS follow the main sequence in both heterogeneous- and homogeneous-discriminability conditions.

### Microsaccades and task performance

We separated the data based on whether the participant made a microsaccade during the critical period for performance (between ready signal onset and stimulus offset) or not. Based on this criterion, the number of trials with a MS (12,789 in heterogeneous-, and 12,051 in homogeneous-conditions) and without (12,583 in heterogeneous-, and 12,402 in homogeneous-conditions) were similar in both conditions across all participants. A three-way ANOVA (2 (experiment 1 vs 2) x 2 (MS presence vs absence) x 4 (locations)) revealed a significant three-way interaction *(F*(15,432) = 53.06, *p < .001)* (**Figure 5a**). For Experiment 1, a two-way ANOVA yielded main effects of location *(F*(7,223) = 81.69, *p < .001)*, showing that both asymmetries are present (RHM, LHM > UVM, LVM, *p < .001*; LVM > UVM, *p < .001*), and microsaccade presence *(F*(7,223) = 17.70, *p < .001)*, but no interaction between them. Performance was better in the absence (1.56±0.05) than presence (1.36±0.07) of microsaccades (**Figure 5b, left panel**). For Experiment 2, a two-way ANOVA yielded only a main effect of microsaccade presence *(F*(7,223) = 20.09, *p < .001)*. Again, performance was better in the absence (1.60±0.05) than presence (1.27±0.07) of microsaccades (**Figure 5b, right panel**) during the critical period.

**Figure 5.**
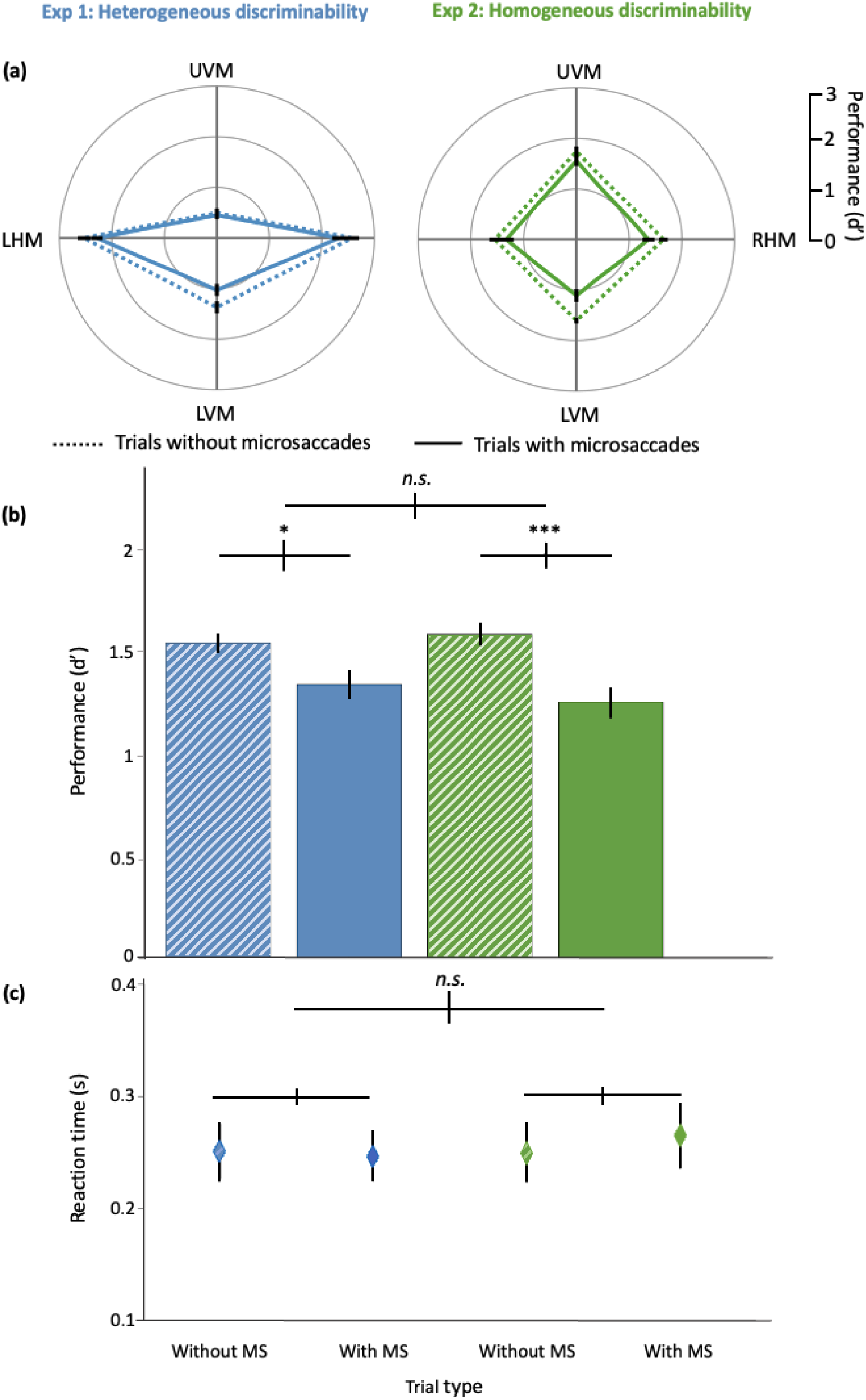
(a) The polar plots show mean d’ performance at all tested locations on trials with (solid line) and without (dotted line) microsaccades, with black lines representing ±1 SEM. (b) Bar plot with group average d’, (c) Plot with group average reaction times on trials with (solid fill) and without (pattern fill) microsaccades during the critical period in both heterogeneous-(blue) and homogeneous-(green) discriminability conditions. Error bars are ±1 SEM. n.s. = not significant; * p < .05; *** p < .001

Regarding RT (**Figure 5c**), neither the interactions nor the main effects were significant (all p > .05). Thus, there were no speed-accuracy tradeoffs.

### Microsaccade rate signature

For both experiments, from the beginning of every trial, MS rate steadily decreased from ∼2 Hz in anticipation of the predictable stimuli onset, decreasing even more rapidly after the onset of the ready signal (which caused the expected transient increase) and in anticipation of the display onset; ‘pre-target inhibition’ was lowest when the stimuli were onscreen (∼0.4 Hz), then rebounded shortly after the onset of the informative response cue (‘post-target rebound’) (**Figure 6**).

**Figure 6.**
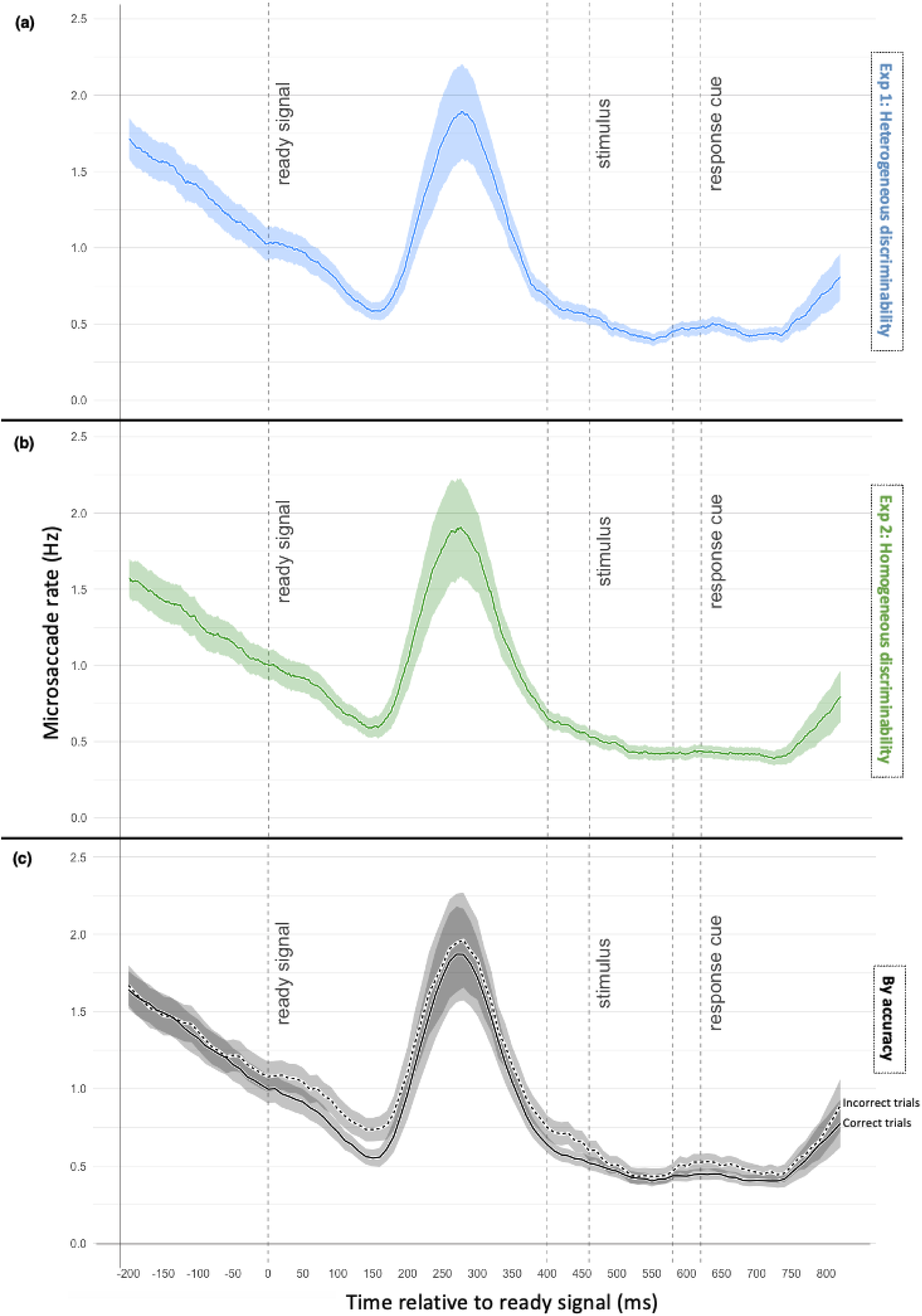
Microsaccade rate timeseries for (a) Experiment 1 (heterogeneous discriminability, blue), (b) Experiment 2 (homogeneous discriminability, green), and (c) sorted by task performance accuracy across both experiments. In panel (c), the solid line represents the temporal dynamics of microsaccades in correct trials; the dotted line, of incorrect trials from both experiments. Cluster-based permutation tests showed no difference both between the two experiments (not depicted) or between correct and incorrect trial data across the experiments.

We compared this MS rate signature for both experiments (**Figure 6a,b**), as well as for correct and incorrect trial data across experiments (**Figure 6c**), using a cluster-based permutation test and found no statistically significant differences (all *p* > 0.5) across the entire trial sequence. We analyzed correct vs incorrect trials across experiments after confirming that they did not differ in each separate experiment.

### Microsaccade direction

The pattern of MS directions across the entire trial sequence was highly similar under both conditions of heterogeneous (Experiment 1) and homogenous (Experiment 2) discriminability (**Figure 7b**). A 2-way ANOVA (2 (experiment 1 vs 2) x 4 (locations) revealed no significant interaction or any main effects (all *p* > 0.05).

**Figure 7.**
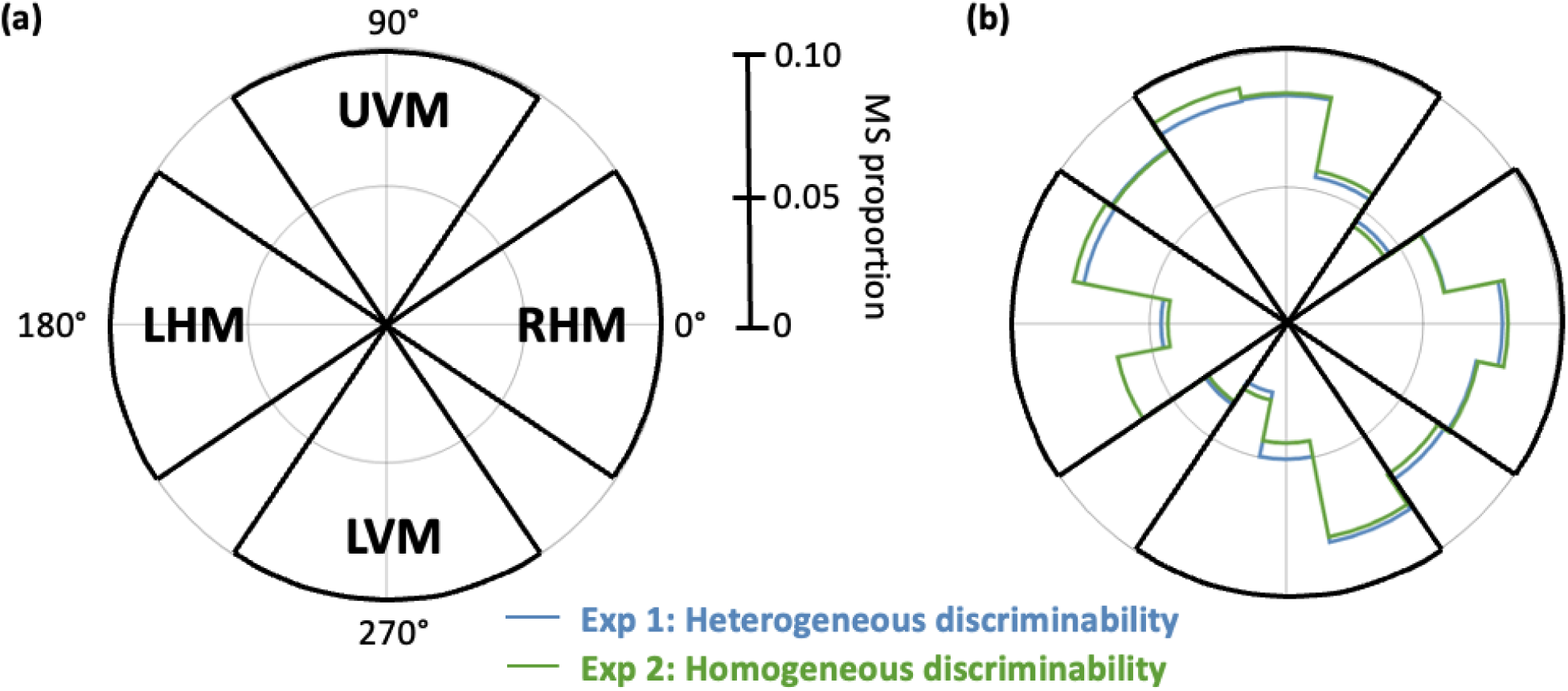
Mean microsaccade direction (a) segments (67.5°, outlined in black) along each cardinal direction used for statistical testing of directional bias, (b) observed over the entire trial and split by experiment.

Given that during the response period observers knew the location where the target had been, we conducted a complementary analysis of MS directions in which we examined response period data separated by target location. A three-way ANOVA (2 (experiment 1 vs 2) x 4 (targets) x 4 (locations)) revealed a significant two-way interaction between target and location (with Greenhouse-Geisser correction) *(F*(2.84,76.66) = 15.45, *p < .001)* (**Figure 8**). A post-hoc analysis of a two-way ANOVA (4 (targets) x 4 (locations)) indicated that a significantly greater number of microsaccades were directed toward the target location during this period with no location uncertainty. This was true regardless of the target location, as well as the discriminability condition.

**Figure 8.**
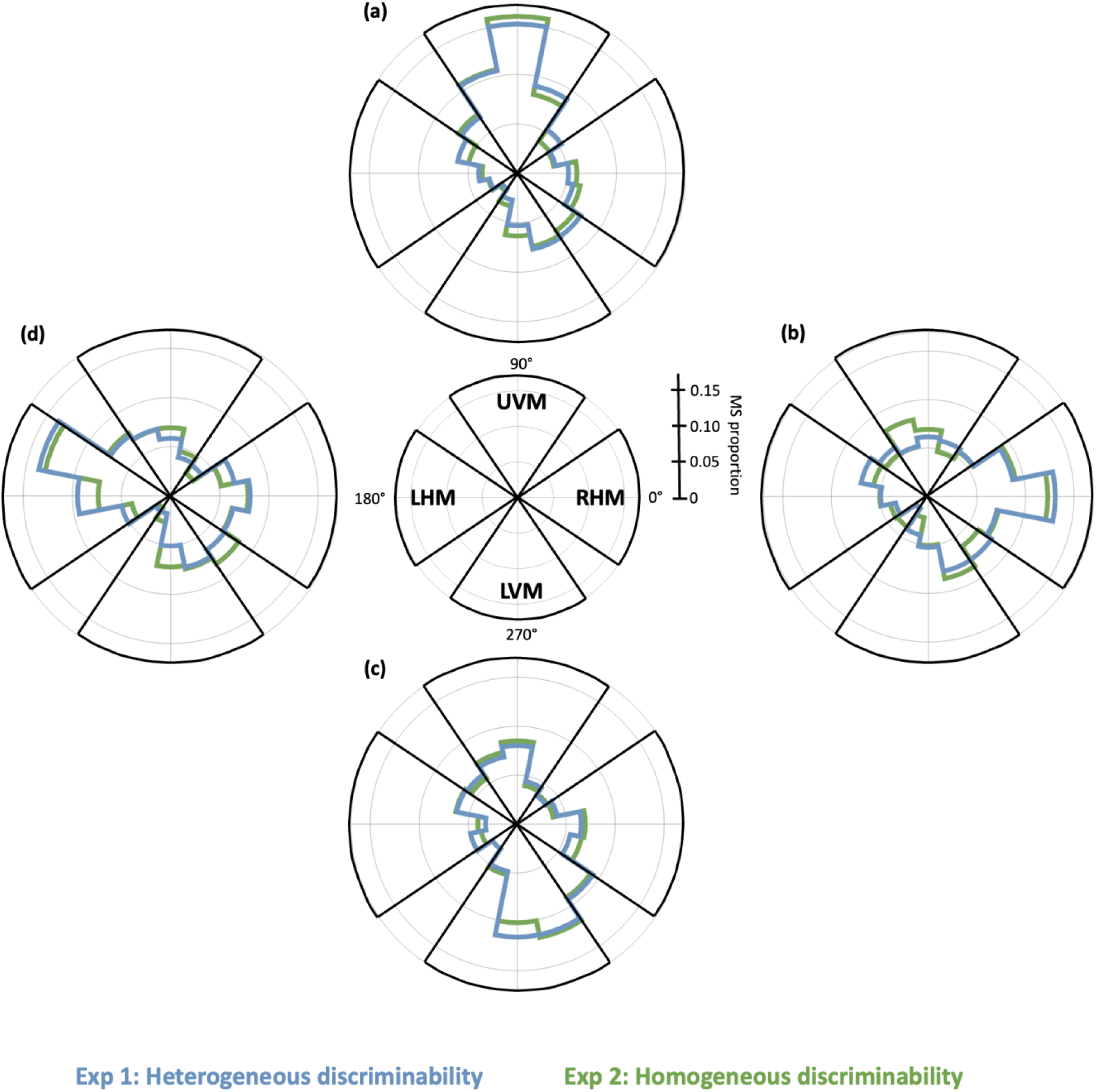
Mean microsaccade direction during the response period when the target was at the (a) UVM, (b) RHM, (c) LVM, and (d) LHM for both experiments. The central polar plot and associated guidelines may be used as the reference scale and orientation for all direction plots in the figure.

### Gaze position

In both experiments, we found that the mean gaze position across observers throughout the trial sequence as well as during the response period was not biased away from the screen center in any direction (within ∼0.2°; **Figure 9**). In all cases it was barely below the horizontal meridian and barely right of the vertical meridian. Thus, we can rule out that corrective microsaccades for fixation as responsible for any of the above reported directional effects.

**Figure 9.**
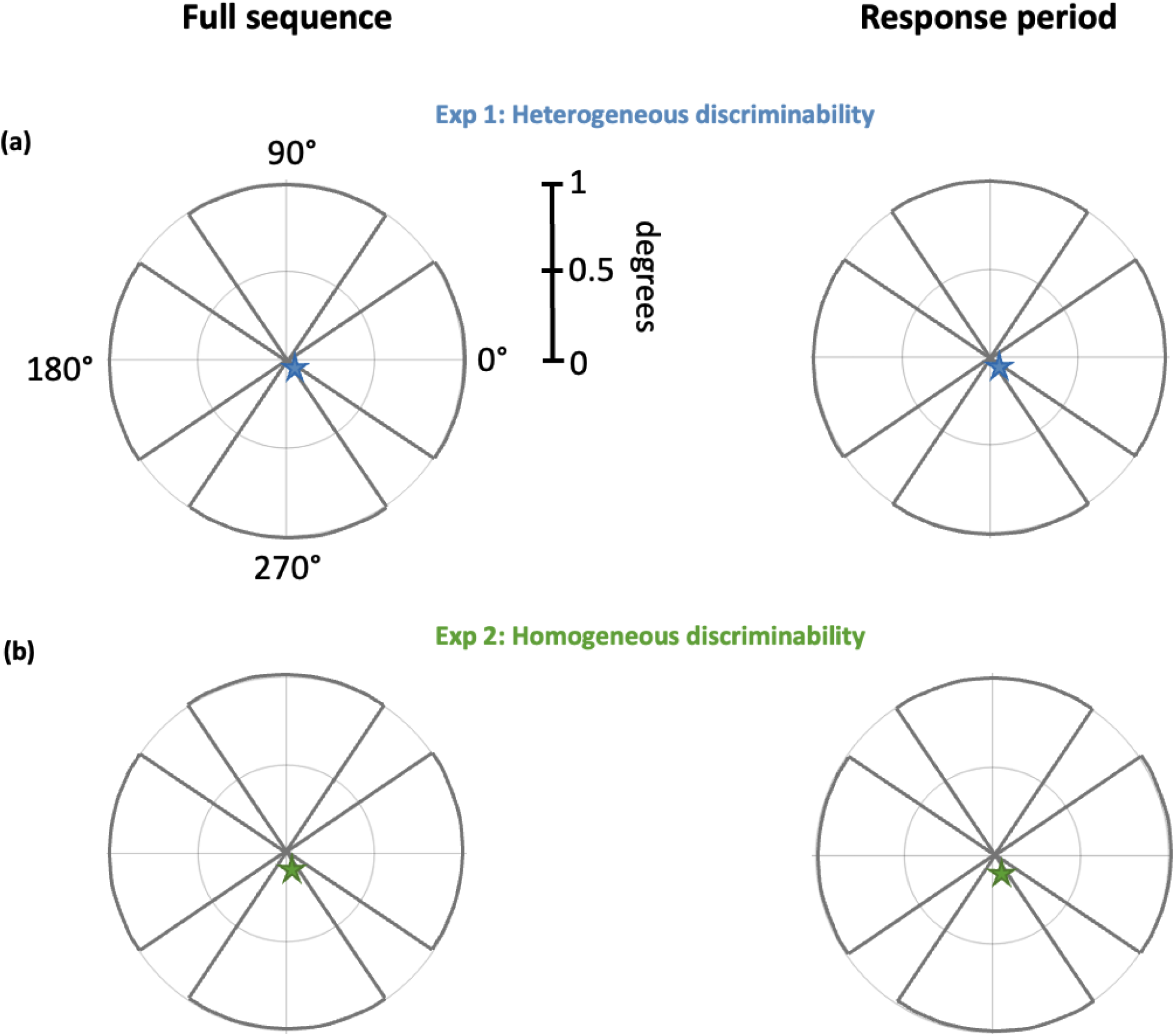
Gaze position (star) across the full sequence (left panels) and response period (right panels) for (a) Experiment 1 (heterogeneous discriminability) and (b) Experiment 2 (homogeneous discriminability). The axis range and angular markers remain constant within all gaze density plots; the full sequence panel (a, left panel) may be used for reference.

## Discussion

We conducted this study to investigate whether and how MS rate and direction are modulated around the visual field (performance fields), whether this pattern changes when discriminability varies or is equated, and whether microsaccades compensate for poor stimulus visibility and its corresponding low discriminability. The results revealed that behavioral performance was better in the absence of MS, and we confirmed that there were no speed-accuracy tradeoffs. The microsaccade temporal and directional patterns were similar under both conditions of heterogeneous and homogeneous discriminability: The microsaccade temporal pattern followed the well-characterized microsaccade rate signature – consistent rate during the precue period, followed by a pre-target inhibition, reaching the lowest rate during stimulus presentation, and culminating in a post-target rebound –, which did not differ according to trial accuracy; there was no significant bias of microsaccades in any direction throughout the trial, except during the response period, when MS were biased toward the location where the target had appeared.

### Microsaccades and task performance

In the present study, we found that that the presence of MS was associated with poorer task performance, regardless of whether stimulus discriminability varied by location (Experiment 1) or was equated (Experiment 2) (**Figure 5b**). Thus, MS did not flexibly adapt to task requirements to help compensate for lower discriminability around the visual field, either in the presented display or where discriminability is typically lower. This could partially be explained by the well-established oculomotor inhibition effect, which describes the steady decline in MS rate before the predictable onset of a brief stimulus (Dankner et al., 2017; Amit et al., 2019; Abeles et al., 2020; Badde et al., 2020; Hung et al., 2023), likely supporting vision by reducing eye movement occurrence during target presentation, which could impair task performance.

Early studies relating performance and eye movements assumed that fixational eye movements aid visual performance (e.g., Averill & Weymouth, 1925; Adler & Fliegelman, 1934). However, whereas these early studies were foundational in showing that the eyes are rarely completely still and move even during fixation, they suffered from methodological limitations, as they had elaborate eye tracking setups with poor sampling resolution. Recent studies with more rigorously timed psychophysical experiment protocols and higher resolution tracking have provided more nuanced insight, suggesting that MS may not be necessary for performance improvements (e.g., Rucci & Desbordes, 2003; van Ede et al., 2019; Liu et al., 2022). Rucci and Desbordes (2003) analyzed the effect of retinal stabilization on discriminating the orientation of a low-contrast and noisy small bar for a brief (500ms) and longer (2s) display duration. Their results showed a greater impairment in correct visual discrimination of briefly presented stimuli (500ms), as opposed to stimuli presented for longer time intervals (2s). They also found that the difference in performance between stabilized and unstabilized conditions is also greater for trials with a longer (2s) stimuli display duration. They speculated that the motion of the retinal image plays a role in refreshing, and possibly structuring, neural activity during brief visual fixation periods. The authors suggested that a longer stimulus presentation period results in an improvement in discrimination performance by allowing both a longer period for affecting the statistical structure of neural activity – as supported by their model – and higher instability of visual fixation.

In a study investigating oculomotor recruitment during a simple working memory task, participants were asked to memorize multiple colored and oriented bars and reproduce the orientation or color of one bar after a short memory delay (van Ede et al., 2019). The researchers found that gaze shifted in the direction of the internal space of memory (i.e., increased number of gaze shifts towards the memorized location of the probed item), even though the spatial locations of the memory items were incidental and not required for task performance. In another study exploring the relation between MS and performance in a covert spatial attention task, participants were asked to report the orientation of a memory item (a tilted bar) based on the color match with an earlier-presented fixation cross (Liu et al., 2022). The authors found that the direction and timing of microsaccadic- and EEG-alpha-modulations of spatial attention were correlated, but trials with no attention-driven microsaccades nevertheless showed preserved alpha modulation by covert spatial attention. Moreover, there were no differences in alpha modulation between trials with or without attention-driven MS. These results suggest that the presence of microsaccades is not a prerequisite for covert spatial attention-modulated neural or behavioral changes.

In addition, other studies (e.g., Horowitz et al., 2007; Roberts & Carrasco, 2019; Li, Pan, & Carrasco, 2021; Palmieri et al., 2023; Hung & Carrasco, 2023; Hung et al., 2023) have shown that presence of MS does not significantly modulate performance. For example, in Hung & Carrasco (2023), the authors found that microsaccade rates during pre- and post-target intervals were similar between correct and incorrect trials. Similarly, in the temporal attention task used in Palmieri et al. (2023), the authors also found that the occurrence of MS did not affect task performance or the magnitude of the temporal attention effect. Overall, this pattern emerges when the criteria for MS presence is if one or more such eye movements is/are detected within a time period that is critical for the experiment (e.g. between the onset of an informative cue and the offset of a stimulus) (e.g., Horowitz et al., 2007; Roberts & Carrasco, 2019; Abeles et al., 2020; Li, Pan, & Carrasco, 2021; Palmieri et al., 2023) as well as when MS are binned into several intervals and analyzed across the trial (Hung et al., 2023). These studies encompass different tasks (e.g., orientation discrimination or Landolt acuity) and phenomena under study, including covert endogenous- and exogeneous-attention, presaccadic attention, feature-based attention, working memory, and perceptual learning. Therefore, the role of MS in modulating performance is complex, and the conditions under which MS presence aids, deters, or has little effect on performance need further systematic investigation.

### Microsaccade dynamics

In agreement with previous studies (Barlow, 1952; Zuber et al., 1965; Bahill et al., 1975), we found that MS amplitudes and peak velocities were highly correlated (**Figure 4**). Because saccades also follow this main sequence (i.e., saccade amplitudes and peak velocities are highly correlated), this finding supports the existence of a ‘microsaccade-saccade continuum’ (Otero-Millan et al., 2008; Otero-Millan, Macknik et al., 2011; Yuval-Greenberg et al., 2014; Rucci & Poletti, 2015; Dankner et al., 2017). Also similar to previous studies, we found that MS rate across the trial sequence follows a reliable pattern (**Figures 6a-c**): decreasing from the beginning of each trial in anticipation of the predictable stimuli onset, and more rapidly after the onset of the ready signal in anticipation of target onset (‘pre-target inhibition’; e.g., Engbert & Kliegl, 2003; Rolfs et al., 2008; Denison et al., 2019; Abeles et al., 2020; Badde et al., 2020; Palmieri et al., 2023), are most inhibited during stimuli presentation, and rebound shortly after the onset of the response cue (‘post-target rebound’). Importantly, these patterns were notably similar between both experiments, regardless of whether stimulus discriminability varied across locations or not (**Figures 6a-b**), and whether the observer had correctly or incorrectly discriminated the target orientation (**Figure 6c**).

### Microsaccade direction

We investigated, for the first time, whether and how the directionality of microsaccades varies when stimuli are simultaneously placed along each cardinal, both under the typical viewing condition of heterogeneous discriminability, as well as under an artificially induced condition of homogeneous discriminability by adjusting stimulus contrast. We found that the proportion of MS toward each direction was the same regardless of discriminability conditions (**Figure 7b**), and even when analyzed by specific trial events. MS proportion was similar towards all locations, except during the response period.

A recent study from our lab (Palmieri et al., 2023) analyzed microsaccade direction in a temporal attention task with one stimulus placed at either the fovea, UVM or RHM. The authors found that microsaccades were directed to the upper hemifield regardless of target location or (neutral vs temporal attention) condition, even though there was no location uncertainty as there was only one stimulus per trial and the target was presented at the same location throughout a whole session. In contrast, in the current study, there was target uncertainty until the response cue appeared, as four stimuli were simultaneously presented and interestingly, at that time we found that the direction of the microsaccades was biased toward the location where the target had been. A similar bias during the response period was found in a recent visual perceptual learning study from our lab, and that tendency increased after training (Hung et al., 2023).

Both the present study and the recent study by Palmieri et al. (2023) are inconsistent with earlier studies that have reported a horizontal bias in MS direction patterns (e.g., Engbert & Kliegl, 2003; Laubrock et al., 2005; Liang et al., 2005; Turatto et al., 2007; Yokoyama et al., 2012; Yuval-Greenberg et al., 2014; Raveendran et al., 2020). Critically, all but two of these studies placed stimuli only to the left and right of fixation, or at fixation: Yuval-Greenberg et al. (2014) also used a 2-AFC orientation discrimination task with stimuli placed at 8 isoeccentric locations around the visual field, rather than the four we used in our study. However, importantly, their stimuli- and cue-presentation was contingent on a MS occurrence and its direction. This is a critical difference from our study, as our design was not dependent on the occurrence of a microsaccade. Even after controlling the retinal eccentricity of the targets, they found a horizontal bias in MS direction, with fewer in the upward direction, and the least in the downward direction. It is likely that spontaneous microsaccades – as in the study by Yuval-Greenberg et al., (2014) – function similarly to spontaneous saccades (e.g., Otero-Millan et al., 2011) and demonstrate a horizontal bias regardless of the number, location, and eccentricity of stimuli. To relate MS occurrence to performance, Yuval-Greenberg and colleagues used a postcue following the appearance of MS- contingent stimuli. They found that performance was better in congruent trials, in which the postcue was in the same direction as the detected microsaccade, than in incongruent trials. In contrast, we found that, on average, the occurrence of a MS during a critical period, regardless of target congruency, was detrimental to performance.

In Turatto et al. (2007), the authors used a feature-search task in a focused versus distributed attention mode. They placed equidistant, colored, cut-off diamonds at a 10° eccentricity and participants had to detect the presence of a singleton target (detection task; their ‘distributed attention’ condition) or had to discriminate which side the target diamond was cut (discrimination task; their ‘focused attention’ condition). They also used a more liberal microsaccade detection criterion (eye movements ≤ 2°) than most MS studies. Given the differences in experimental conditions and analysis, our results are not directly comparable. However, it is worth noting that they did not find a significant directional bias in any direction in the detection.

### Microsaccade direction: Response period

When analyzing the response period data – once the stimuli were no longer present and there was no longer uncertainty regarding the target location – we found that the microsaccade direction pattern under both discriminability conditions was the same (**Figures 8a-d**): A significantly greater number of microsaccades were directed toward the target. This finding is consistent with studies reporting a MS directional bias towards the target location in perceptual learning (Hung et al., 2023) and working memory (van Ede et al., 2019) tasks, once the stimuli were no longer present and an informative response cue was presented indicating the target location.

These directional biases are likely an automatic orienting of microsaccades toward the location of greatest interest (the target) during the response period. They might reflect an attempt by the memory-guided oculomotor system (Meister & Buffalo, 2016; van Ede et al., 2019) to retrieve target information indicated by the response cue before preparing a response (Willeke et al., 2019). This idea is supported by results from a recent study in which the authors studied MS direction patterns in the macaque monkey during a visual discrimination task (Raffi et al., 2021). In the trial sequence, a spatially localized cue was presented after the appearance of an optic flow stimulus, signaling the beginning of the response period (release of a lever within time constraints). During this time period, as well as other analyzed epochs, they found that MS directions were significantly clustered toward the visual cue. All four studies, including the present one, suggest that MS proportion is biased in the direction of the target during the period without location uncertainty, and could indicate an attempt to retrieve target characteristics, even though the stimulus is no longer present.

Another study has reported different findings: In their discrimination (‘focused attention’) task, Turatto et al. (2007) found that MS were biased in the direction opposite the target in the 500– 750ms time window, which corresponds to the response period in our study. The authors speculated that the bias was the result of efforts to counteract the execution of a programmed saccade in the direction of the target towards which attention was necessarily shifted to perform the task. However, given that their task sequence did not include a ready signal and/or a response cue, their analysis period was not tightly locked. Earlier studies have reported MS directed towards the cued location immediately after an attention cue and then away from the cued location (300ms) after stimulus onset (Rolfs et al., 2004; Laubrock et al., 2005). Both these studies used attentional cues or imperative stimulus onsets, but because they did not use distinct response cues, a response period cannot be isolated. Together, these findings suggest that microsaccade directionality is not driven by the same factors throughout the trial sequence.

### Conclusion

In this study, we used a simple design (2AFC task, stimuli placed along the cardinal directions) and optimized trial event timing to explore whether and how MS and behavioral performance around the visual field are linked. We found that microsaccade rate and directionality do not vary with differing discriminability. The MS temporal pattern followed the typical ‘microsaccade rate signature’ and was the same for both conditions of heterogeneous and homogeneous discriminability. In examining MS directionality, we found that it followed neither the performance field pattern found in behavior, nor the horizontal bias reported in microsaccade literature. Placing stimuli at cardinal locations, we found no overall bias in MS proportion toward any direction. Once observers knew the location where the target had been, microsaccades were biased in that direction. Importantly, we found that the presence of MS during the critical stimulus period (between ready signal onset and stimulus offset) lowered performance. Overall, we found that microsaccade patterns (temporal and spatial) are similar with varying – heterogeneous or homogeneous – discriminability. Together, the results indicate that MS do not flexibly adapt to task requirements to compensate for lower discriminability around the visual field. This novel finding is interesting given the resilient nature of PFs. Several studies have examined PFs and their retinal and cortical correlates (review: Himmelberg et al., 2023), and the findings of the present and a recent lab study (Palmieri et al., 2023) highlight the importance of including microsaccades as an oculomotor factor in studying these visual asymmetries.

This study brings the field a step closer to assessing eye movements in naturalistic settings, because we are more likely to encounter objects with different degrees of visibility around our visual field than only along the horizontal meridian, for example. They have the potential to inform models of MS performance and oculomotor function in general, given that MS and saccades are believed to share performance characteristics and the same neural generator. In addition, given how some MS characteristics differ in atypical and visually impaired populations (e.g., Otero-Millan et al., 2013; Dankner et al., 2017; Panagiotidi et al., 2017; Alexander et al., 2018), future research could examine whether the MS pattern we observed differs in such populations and could be used as a non-invasive diagnostic indicator in neurological diseases.

## Acknowledgments

This research was supported by NIH-National Eye Institute RO1-EY027401 to M.C. The authors would like to thank Carrasco Lab members – Aysun Duyar, Rania Ezzo, Shao-Chin Hung, and Helena Palmieri, in particular – for their valuable feedback.

